# Selective deletion of *Tsc1* from mouse cerebellar Purkinje neurons drives sex-specific behavioral impairments linked to autism

**DOI:** 10.1101/2024.08.07.607071

**Authors:** Ryan J. Lawson, Nicholas J. Lipovsek, Samuel P. Brown, Achintya K. Jena, Joanna J. Osko, Joseph L. Ransdell

**Affiliations:** Department of Biology, Miami University, Oxford, OH 45056

**Keywords:** tuberous sclerosis, Tsc, cerebellum, social interaction, motor coordination, sexual dimorphism

## Abstract

There is a striking sex bias in the prevalence and severity of autism spectrum disorder (ASD) with 80% of diagnoses occurring in males. Because the molecular etiology of ASD is likely combinatorial, including interactions across multiple genetic and environmental factors, it is difficult to investigate the physiological mechanisms driving sex-specific differences. Loss of function mutations in *TSC1* result in dysregulated mTORC1 signaling and underlie a multi-system disorder known as tuberous sclerosis (TSC). Interestingly, more than 50% of individuals diagnosed with TSC are also diagnosed with ASD, making TSC mutations one of the most prevalent monogenic causes of ASD. Mice harboring targeted deletion of *Tsc1* selectively in cerebellar Purkinje neurons, referred to here as *Tsc1^mut/mut^*, have multiple ASD-linked behavioral impairments, including deficits in social interactions, motor coordination, and vocalizations. However, these ASD-linked behavioral deficits have only been investigated using male *Tsc1^mut/mut^* animals. Here, we used cohorts of male and female *Tsc1^mut/mut^* animals to determine if behavioral impairments, previously identified in this model, are similar across sex. Specifically, we measured balance and motor coordination and social interaction behaviors in two age groups across sex. W*e* determined balance and motor coordination deficits are similar in male and female *Tsc1^mut/mut^* mice, and that deficits in the firing of *Tsc1^mut/mut^* Purkinje neurons located in the cerebellar vermis are also similar across sex. However, impairments in social approach behavior were found to be significantly more severe in *Tsc1^mut/mut^* males compared to females. These results indicate the selective deletion of *Tsc1* in Purkinje neurons differentially impairs cerebellar circuits based on sex.

## 1. INTRODUCTION

Across individuals, measures of ASD symptom manifestation and severity are heterogeneous (Geschwind 2011), however, a highly replicated finding in ASD research is the greater number of diagnoses in males compared to females (Christensen et al. 2018). Sex-specific ASD differences have also been described but vary across age groups and symptom domains (Van Wijngaarden-Cremers et al. 2014; Tillmann et al. 2018; Craig et al. 2020). While the details of these sex-specific differences continue to be revealed, it remains difficult to investigate the underlying cellular and molecular mechanisms due to the etiology of ASD typically involving multiple interacting genetic and environmental factors. Here we examine if there are sex specific phenotypes in an ASD-mouse model which has selective deletion of a single ASD-linked gene, *Tsc1*, from a single neuronal cell type, cerebellar Purkinje neurons.

Several lines of evidence implicate cerebellar dysfunction in the development of ASD (Wang, Kloth, and Badura 2014). Cerebellar Purkinje neurons integrate incoming information and are the sole output cells of the cerebellar cortex (Palkovits, Magyar, and Szentágothai 1972; Andersen, Korbo, and Pakkenberg 1992). Reduced Purkinje neuron firing has been directly linked to ASD-like behavioral deficits in mice (Tsai et al. 2012; 2018; Kalume et al. 2007) and post-mortem investigations have revealed a reduced number and density of cerebellar Purkinje cells in individuals with ASD (Wang, Kloth, and Badura 2014; Bailey et al. 1998; Stoodley 2014). Tuberous sclerosis complex (TSC) is a disorder affecting multiple organ systems with severe cognitive, behavioral, and psychiatric symptoms (Crino, Nathanson, and Henske 2006). TSC is the result of loss-of-function mutations in TSC1 or TSC2 (European Chromosome 16 Tuberous Sclerosis Consortium 1993; Au et al. 2007). Proteins encoded by these genes are necessary to form a signaling complex which drives inhibition of the mammalian target of the rapamycin complex 1 (mTORC1) pathway, which plays a major role in regulating cell growth and metabolism (Wullschleger, Loewith, and Hall 2006). Interestingly, greater than 50% of individuals diagnosed with TSC also meet diagnostic criteria for ASD (Wiznitzer 2004; Jeste et al. 2008), making TSC mutations one of the most prevalent monogenic causes of ASD.

To evaluate the role of Tsc1 in cerebellar PNs, Tsai et al., (2012) developed a mouse model with targeted and selective *Tsc1* deletion in cerebellar Purkinje neurons using L7-Pcp2:Cre/LoxP recombination (Barski, Dethleffsen, and Meyer 2000; Saito et al. 2005). Interestingly, this mouse model, referred to here as *Tsc1^mut/mut^*, exhibits several ASD-linked behavioral phenotypes, including deficits in motor coordination, vocalizations, social interactions, and exaggerated repetitive behaviors (Tsai et al., 2012). These behavioral impairments have been linked to both reduced repetitive firing in cerebellar Purkinje neurons, and in older animals (>10 weeks-old), to the progressive loss of Purkinje neurons via apoptosis (Tsai et al., 2012; 2018). These previous investigations of *Tsc1^mut/mut^* phenotypes, however, did not include female test subjects.

We examined if *Tsc1* deletion differentially affects Purkinje neuron intrinsic excitability and ASD-linked behavioral impairments across male and female adult mice. We determined that in older (16-24 week-old) *Tsc1^mut/mut^* mice, social interaction deficits are significantly more severe in males compared to females, however, balance and motor coordination impairments in these animals are similar across sex, suggesting cerebellar circuits are differentially impaired in *Tsc1^mut/mut^*mice based on animal sex.

## 2. METHODS

### 2.1 Subjects

This study used the L7-*Pcp2*:Cre/*Tsc1* floxed animal model of autism in mice (*Mus musculus)* with a C57BL/J6 genetic background (Jackson Laboratories). After weaning, mice were housed with littermates of the same sex at Miami University’s Laboratory of Animal Resources facility, with controlled temperature and humidity and a 12:12 light cycle starting at 7:00am. Mice were provided with water and a standard rodent diet ad libitum (Lab Diet 5001, Cincinnati Lab Supply Inc., Cincinnati, OH, USA). All animal experimentation was approved by the Institutional Animal Care and Use Committee of Miami University (protocol #1044). Cre-negative *Tsc1*^flox/flox^ mice were crossed with Cre-positive *Tsc1*^flox/wt^ mice to generate Cre-positive *Tsc1*^flox/wt^, referred to here as *Tsc1*^mut/wt^, and Cre-positive *Tsc1*^flox/flox^ mice, referred to here as *Tsc1*^mut/mut^ mice for experiments. These pairings also generated Cre-negative *Tsc1^flox/flox^*mice, which were used as littermate controls in behavioral studies. L7-Pcp2 Cre-positive animals (lacking loxP sites) and Cre-negative *Tsc1^flox/flox^*have previously been shown to not differ from wild type mice in behavioral assays (Barski, Dethleffsen, and Meyer 2000; Ehninger et al. 2008; Tsai et al. 2012; 2018).

### 2.2 Microscopy and image processing

Parasagittal cerebellar slices (25 µm) of fixed and cryoprotected brains were used for imaging the tdTomato fluorescent signal in Cre-positive Purkinje neurons. Images were acquired using a Zeiss LSM-710 confocal microscope using either a 63X oil emersion objective or 20X objective. Z-series stack images with 1 µm intervals or single plane snapshots were used to assess tdTomato expression in Cre-reporter mice.

### 2.3 Electrophysiology Methods

Spontaneous and evoked action potentials were recorded from Purkinje neurons in parasagittal brain sections acutely isolated from 5-8 week-old animals. Brain sections were perfused with warmed (33-34°C) artificial cerebral spinal fluid (ACSF) that was continuously bubbled with 95% O2/5% CO2. Borosilicate glass recording pipettes were filled with an internal solution containing (in mM): 0.2 EGTA, 3 MgCl2, 10 HEPES, 8 NaCl, 4 Mg-ATP, and 0.5 Na-GTP. Recording pipette resistances were 2–4 MΩ. Electrophysiology records were acquired using a dPatch amplifier and SutterPatch acquisition software (Sutter Instrument). Evoked action potential recordings were taken following gap-free spontaneous firing recordings. For evoked firing protocols, cells were initially injected with a −0.5 nA hyperpolarizing current for 100 ms before stepping current injections to 0, 0.1, 0.2, and 0.3 nA for 700 ms. Input resistance and capacitance measurements were obtained from a whole-cell voltage-clamp recording in which membrane voltage was stepped from a −80 mV holding potential to −90 mV (for 100ms) and to - 70 mV (for 100ms) in a subsequent sweep. Membrane capacitance was calculated by integrating the area of fast capacitive transient current during voltage steps and dividing by the change in membrane voltage (10 mV). Electrophysiology recordings were analyzed using SutterPatch (Sutter Instrument), Microsoft Excel (Microsoft), and Prism (GraphPad) software.

### 2.4 Behavioral Tests

Behavioral tests were conducted on male and female mice in cohorts of animals that were 9-11 weeks of age and repeated on animals in the *Tsc1*^mut/mut^ and control cohorts at 16-24 weeks of age. All tests were performed during the light cycle between 9:00 and 17:00.

#### 2.4.1 Elevated Balance Beam

Motor coordination and balance were tested using the elevated balance beam previously described in Carter et al. (2001). The balance beam apparatus consists of an 80 cm-long flat beam 10 mm in width. A second flat beam, 8 mm in width, was also used in this assay and results from the 8 mm beam testing are reported in supplemental Figure 2. During testing, animals traversed the narrow beam to reach an enclosed goal platform which was covered by an opaque plastic goal chamber. As an incentive, food was placed inside the goal chamber. Mice were trained to cross each beam for three consecutive days prior to testing. Each training session included two minutes in which each animal explored the initial starting area of the beam, two minutes to explore the goal chamber, and a single crossing of each (10mm and 8mm) balance beam. After training, mice were tested individually on four consecutive days. Crossing the beam was timed from the moment the animal’s hind limbs crossed the starting line (80 cm from the goal chamber) until the animal’s hind limbs crossed entrance of the goal chamber. The time to cross the beam and the number of hindlimb foot-slips while crossing the beam were recorded and used to assess differences in fine motor coordination and balance across sex and genotype.

#### 2.4.2 Social Behavior

Mice were tested for deficits in social interactions using the three-chamber social interaction task apparatus (Yang, Silverman, and Crawley 2011). The three-chamber apparatus consists of three openly connected chambers constructed of clear plexiglass. Each chamber is 20 cm by 40 cm and has 22 cm high (plexiglass) walls. Each of the two exterior chambers contained an inverted wire cup capable of holding an adult mouse comfortably. A plastic cup was placed on top of the wire cup to prevent climbing (see Fig. 4A). Mice were allowed to acclimate to the testing room for 30 minutes in their home cage prior to testing. The test mouse was placed in the center chamber and allowed to explore the center chamber for seven minutes to acclimate to the chamber environment. The assays were then conducted in three phases, each ten minutes in length: an adaptation phase during which the mouse was allowed to freely roam all three chambers; a social approach phase during which one exterior chamber contained an empty wire cup and the other exterior chamber contained a cup holding an unfamiliar ‘stranger’ mouse (Fig. 4A); and a social novelty phase during which one exterior chamber contained the ‘stranger’ mouse from Session I (now considered familiar) and the previously empty chamber contained a new, unfamiliar stranger mouse (Fig. 6A). ‘Stranger’ mice were animals at least 6 weeks old and within two months of the age of the test mouse, the same sex as the test mouse, and Cre-negative. These animals underwent two 15-minute training sessions prior to being used in testing as a ‘stranger’ to observe for any potentially disruptive behaviors. In all three sessions, the test mouse’s number of entries into each chamber, marked by the full crossing of the hind legs into the chamber, and the time spent in each chamber were scored by an observer with a stopwatch. The total number of direct contacts and the total time in direct contact with the unfamiliar object, unfamiliar mouse, and familiar mouse were also scored by the experimenter using a video recording of the session. Direct contacts were defined as instances of one second or more of direct sniffing or paw-touching of the wire cage. The apparatus and both wire cages were cleaned with 70% ethanol and water between test subjects.

### 2.5 Statistical Analysis

Statistical analyses were completed using Graphpad Prism 8 software (GraphPad Software Inc., La Jolla, CA, USA). Data from whole cell electrophysiology recording measurements were compared across sexes and genotypes using unpaired Student’s t-tests. Data from elevated balance beam experiments were compared across sexes and genotypes using repeated measures (RM) two-way ANOVA tests. Social interaction experiment results were assessed using two-way ANOVA tests; comparisons of the percent of time investigating the unfamiliar animal used sex and genotype as between-subject factors, and comparisons of the number of direct contacts used sex and chamber preference as between-subject factors within each genotype. Šídák’s corrections were applied in multiple comparisons tests. P <0.05 was considered to be statistically significant. P-values for all statistical tests comparing behavioral performance measures across sex are reported in Supplemental Table 1 for the 9-11-week animal cohorts, and in Supplemental Table 2 for the 16-24-week-old animal cohorts.

## 3. RESULTS

### 3.1 *Tsc1* deletion in Purkinje neurons attenuates repetitive firing

A Cre-*LoxP* recombination strategy was used to selectively delete *Tsc1* from mouse cerebellar Purkinje neurons. *Tsc1* mutant mice with *loxP* sites flanking exons 17 and 18 of the *Tsc1* gene (Kwiatkowski et al. 2002) (Jackson laboratory, strain # 005680) were crossed with a transgenic mouse strain that expresses Cre recombinase under the control of the mouse Purkinje cell-specific L7 promoter (Jackson laboratory, strain # 010536) (Fig. 1A). Progeny from this cross included hemizygous Cre-positive animals with either heterozygous or homozygous *Tsc1* floxed alleles, used as test group animals, and Cre-negative animals. Cre-negative animals with homozygous *Tsc1* floxed alleles were used as control group animals. To verify selective Cre expression in cerebellar Purkinje neurons, hemizygous Cre-positive animals were also crossed with the Ai14 Cre reporter tool strain (Jackson Laboratory, strain # 007914) (Fig. 1A), which express tdTomato fluorescence following Cre-mediated recombination (Madisen et al. 2010). Parasagittal cerebellar sections from the Cre-positive progeny of this cross revealed robust tdTomato expression that was selective to Purkinje neurons in the cerebellum of adult animals (Fig. 1B). Cre-expressing Purkinje neurons were identified in male and female animals as young as 11 days old (Fig. 1B). In these sections, no tdTomato fluorescence was identified in cells outside the Purkinje neuron layer.

**Figure 1.**
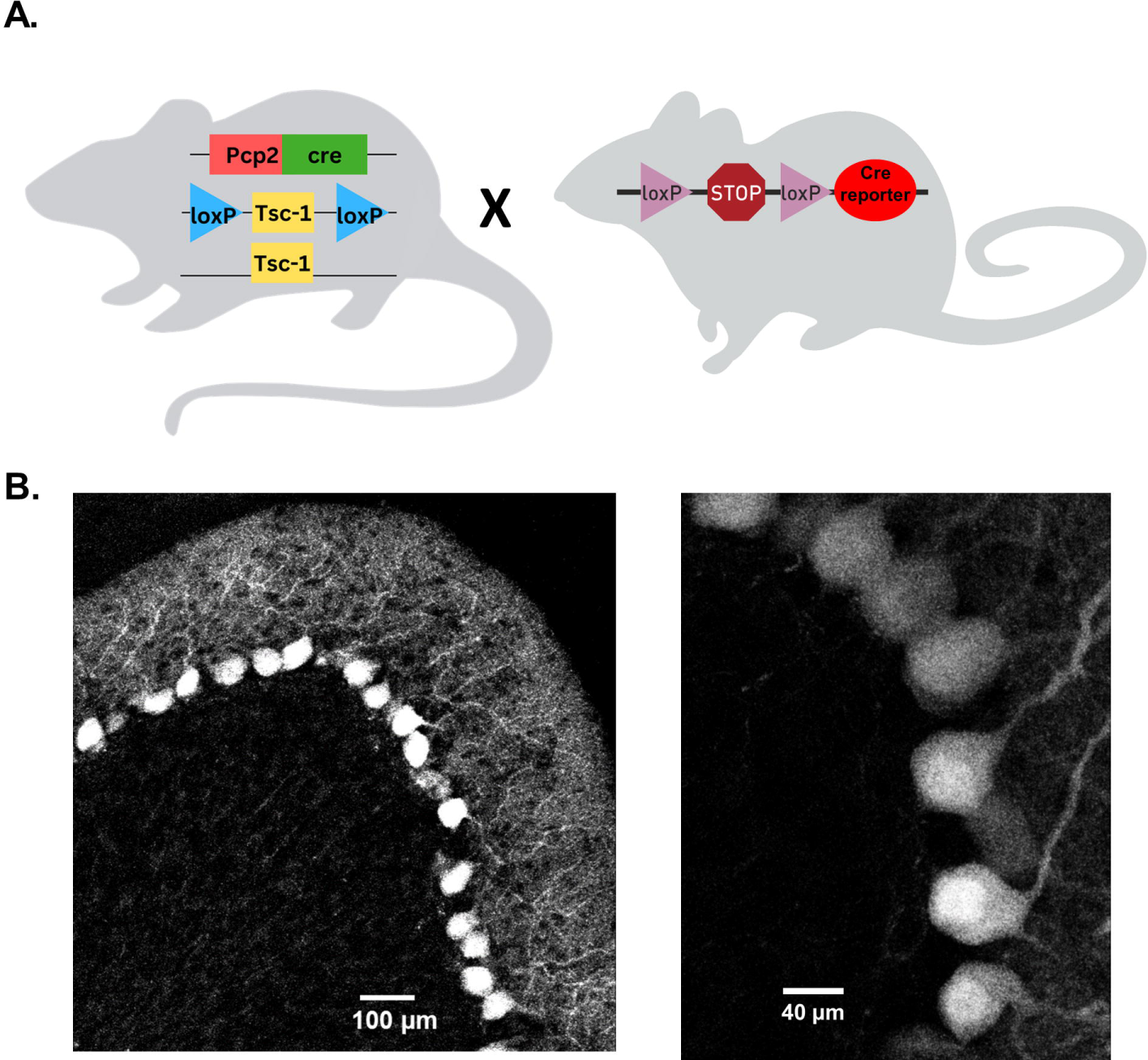
Cre recombinase expression on the L7 Pcp2 promoter can drive selective *Tsc1* deletion in mouse cerebellar Purkinje neurons. Panel **A.** depicts the breeding strategy for Purkinje neuron-specific *Tsc1* deletion. Heterozygous or homozygous animals with *loxP* sites flanking *Tsc1* were crossed with mice hemizygous for the *Cre* recombinase gene under control of a Pcp2 (L7) promoter. To verify the Purkinje neuron-specific expression of Cre recombinase, *Cre-*positive animals were crossed with a Cre reporter strain in which a loxP-flanked STOP cassette prevents expression of a CAG promoter-driven tdTomato gene. In panel **B.**, 20x (left) and 63x (right) confocal images of 25 µm parasagittal cerebellar sections were taken from adult tdTomato Cre-reporter mice expressing Cre on the L7/Pcp2 promotor.

Targeted deletion of *Tsc1* from male (mouse) cerebellar Purkinje neurons has previously been shown to result in attenuated repetitive firing in adult Purkinje neurons (Tsai et al. 2012). To determine if Cre-positive *Tsc1^flox/flox^* progeny (described above), referred to here as *Tsc1^mut/mut^*, have similar deficits in repetitive firing, and to test if changes in membrane excitability is conserved across *Tsc1^mut/mut^* Purkinje neurons from male and female animals, we performed whole cell patch current-clamp recordings from adult (5-8 week-old) *Tsc1^mut/mut^* and wild type Purkinje neurons in acutely isolated cerebellar sections (see Methods). These recordings were performed at near physiological temperatures (33-34°C) and revealed the spontaneous repetitive firing frequency in *Tsc1^mut/mut^* Purkinje neuron is significantly (P<.001, Student’s t-test) reduced compared to age-matched wild type cells (see Fig. 2A, B). Reduced repetitive firing frequencies measured in *Tsc1^mut/mut^* Purkinje neurons were not, however, significantly different between cells isolated from male and female mice (Fig 2C). Depolarizing current injections after a common hyperpolarizing (−0.5 nA) current step revealed similar impaired evoked firing in *Tsc1^mut/mut^* Purkinje neurons, and no difference between male and female *Tsc1^mut/mut^* Purkinje neurons (Fig. 2D, E). Measures of membrane capacitance were also not significantly different across male and female *Tsc1^mut/mut^* Purkinje neurons, however, membrane input resistance was found to be significantly (P=.02, Student’s t-test) lower in male *Tsc1^mut/mut^* Purkinje neurons compared to cells from *Tsc1^mut/mut^* females (Fig. 2F). No sex-specific differences in capacitance and input resistance were measured in wild type controls cells (data not shown).

**Figure 2.**
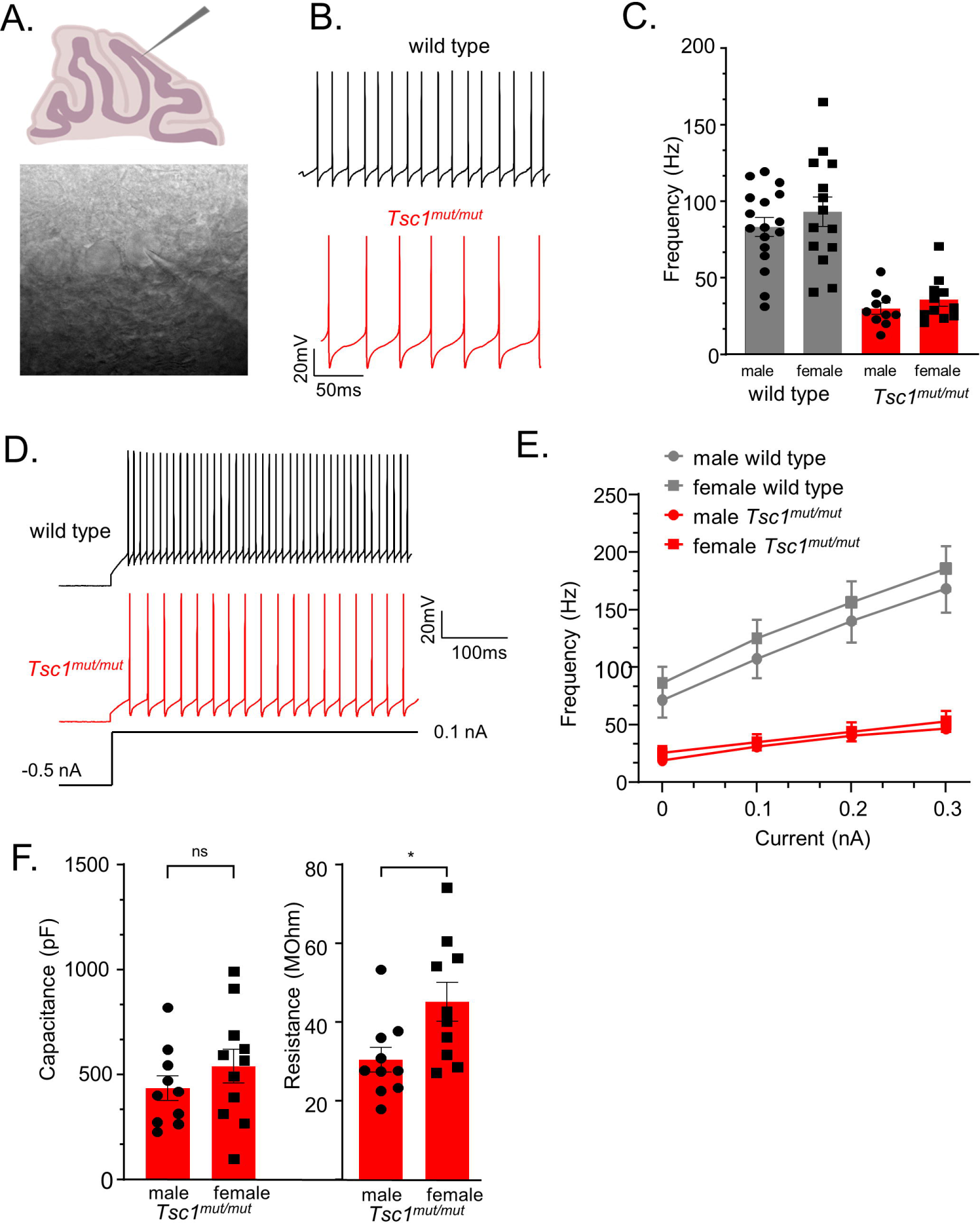
*Tsc1* deletion results in reduced membrane excitability in Purkinje neurons from male and female mice. **A.** 350 µm parasagittal cerebellar sections were acutely isolated for whole cell patch-clamp recordings. In panel **B.**, representative simple spike activity is presented from wild type and *Tsc1^mu/mut^* adult Purkinje neurons. **C.** In *Tsc1^mut/mut^* Purkinje neurons, mean ± SEM spontaneous firing frequency is significantly lower than wild type Purkinje neurons. Within each genotype, mean ± SEM firing frequency is not significantly different across sex. Evoked firing was measured during 0.1-0.3 nA depolarizing current injections after a common −0.5 nA hyperpolarizing step (**D.**). **E.** Compared to wild type controls, mean ± SEM evoked firing frequencies were significantly lower in *Tsc1^mut/mut^* cells, and not different across male and female cells from either genotype. **F.** Mean ± SEM measures of capacitance were not significantly different across male and female *Tsc1^mut/mut^* cells. Mean input resistance was determined to be significantly lower in male *Tsc1^mut/mut^* Purkinje neurons. Input resistance and capacitance did not differ between wild type male and female Purkinje neurons (data not shown).

### 3.2 *Tsc1* deletion from Purkinje neurons impairs motor coordination similarly in males and females

To examine potential sex-specific differences in motor coordination and balance resulting from the targeted deletion of *Tsc1* in Purkinje neurons, the elevated balance beam behavioral assay (Carter et al., 2001) was performed on 9-11 week-old and 16-24 week-old mice. For each age-group, the balance beam test was performed across four consecutive testing days. The amount of time it took the animal to traverse a 10 mm wide beam (see Figure 3A), and the number of hindlimb foot-slips while traversing the beam (see Methods) were recorded. The mean (±SEM) time to cross and hindlimb foot-slips are plotted in Figure 3 for each of the four testing days. Results for the testing of 9-11 week-old animals are plotted in Figures 3B and C, and again for the 16-24 week-old age group in Figures 3D and E. These data revealed that at both ages (9-11-week-old and 16-24 week-old) *Tsc1^mut/mut^* animals have impaired motor performance compared to control (*Tsc1^flox/flox^*; Cre −) mice, taking significantly (Figure 3B, D; P<0.0001, RM two-way ANOVA) longer to cross the 10mm balance beam and having significantly (Figure 3C, E; P<0.0001, RM two-way ANOVA) more hindlimb footslips while traversing the 10 mm beam. Animals with targeted deletion of a single *Tsc1* allele (*Tsc1^mut/wt^*) performed similarly to control animals (P=0.9866, RM two-way ANOVA, Figure 3B). For all genotypes tested on the 10 mm balance beam, no significant differences were recorded across sex (within a genotype). These data suggest there are no significant differences between sexes in ASD-associated deficits in fine motor coordination and balance. This testing paradigm was also performed using a more challenging, 8 mm balance beam, and similar to the 10 mm balance beam, revealed no significant differences across sex within a given genotype (see Supplemental Fig. S1 and S2).

**Figure 3:**
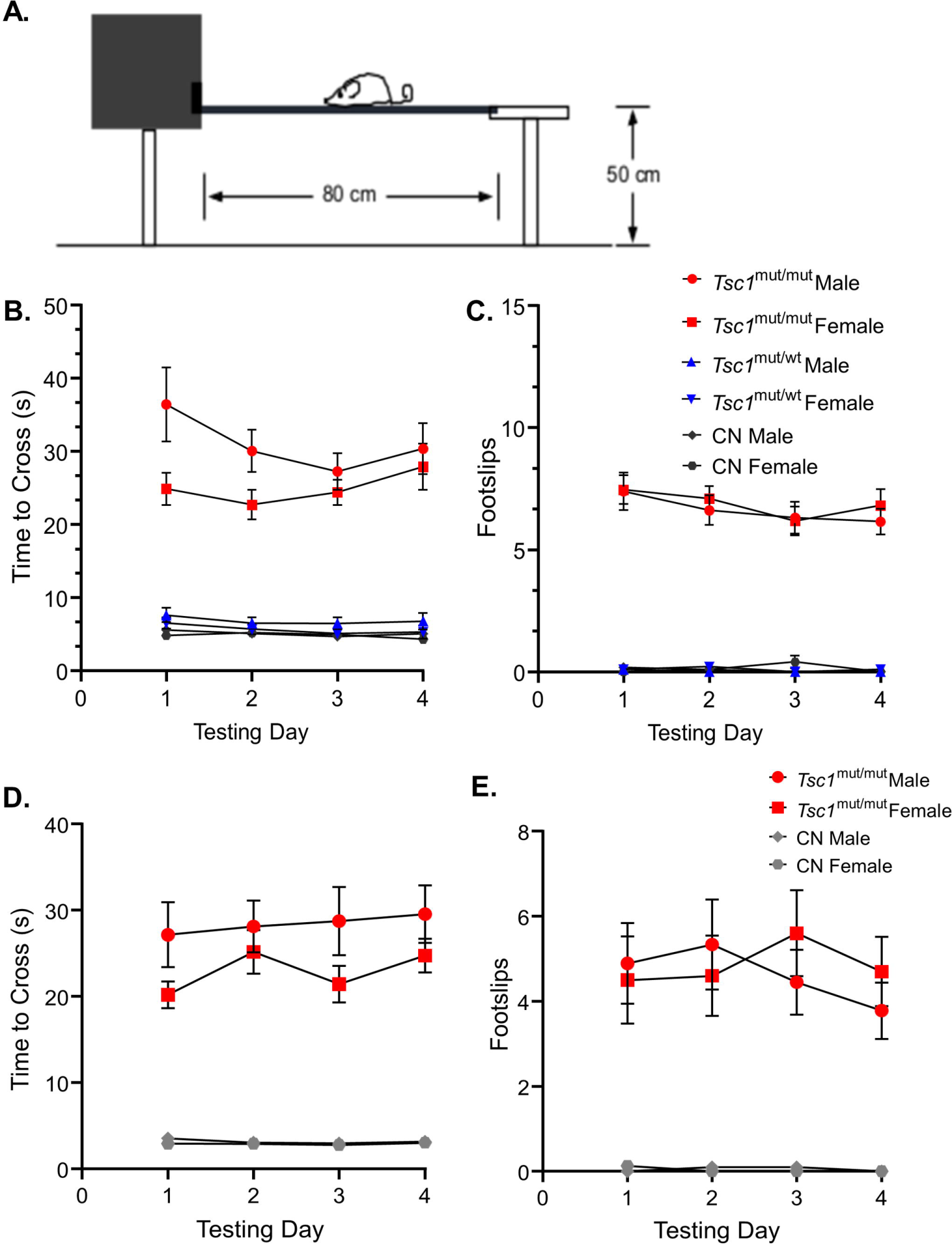
No sex-specific differences were found in performance on the 10mm elevated balance beam in 9-11 week-old and 16-24 week-old mice. **A.** A cartoon depiction of the elevated balance beam testing setup is shown. **B.** 9–11-week-old *Tsc1*^mut/mut^ (N=13 males and N=11 females), *Tsc1*^mut/wt^ (N=10 males and N=10 females), and control (CN, N=11 males and N=10 females) mice were tested on a 10 mm (width) elevated balance beam. Across the four testing days, the mean time to cross of *Tsc1*^mut/mut^ mice (of both sexes) took significantly (P<0.0001, RM two-way ANOVA) longer to cross than *Tsc1*^mut/wt^ and control mice. **C.** The same was true for the number of hindlimb footslips (P<0.0001, RM two-way ANOVA). *Tsc1*^mut/wt^ and control cohorts performed similarly. Within each genotype, male and female animals performed similarly across the four testing days. Mean ± SEM values are plotted. Mean ± SEM results are plotted for16-24-week-old *Tsc1*^mut/mut^ (N=9 males and N=10 females) and control (N=10 males and N=8 females) mice on the 10 mm (width) elevated balance beam in D-E. Within each genotype, male and female animals performed similarly across the four testing days. **D.** *Tsc1*^mut/mut^ mice took significantly longer to cross (P<0.001, RM two-way ANOVA) than CN mice across testing days. **E.** *Tsc1*^mut/mut^ mice had significantly more hindlimb footslips (P<0.001, RM two-way ANOVA) than CN mice across testing days.

### 3.3 Social approach behavior is differentially affected in male and female *Tsc1^mut/mut^* mice

A three-chamber apparatus was used to measure differences between sexes in impairments for social approach preference and social novelty preference in *Tsc1^mut/mut^* mice. The three-chamber apparatus consisted of an empty center chamber and two exterior chambers with wire cups large enough to comfortably hold an adult mouse. In this assay, preference for social approach is reflected as an increase in the investigation of an unfamiliar animal (in one of the wire cups) compared to the alternate empty wire cup (Fig. 4A). In figure 4B, the mean (±SEM) percentage values for the time spent investigating the unfamiliar mouse, as a percentage of the total time spent investigating the unfamiliar mouse and the empty wire cup, are plotted across sex and genotype for 9-11 week-old cohorts. Results from these tests reveal *Tsc1*^mut/mut^ mice spend a significantly (P<0.05, two-way ANOVA) lesser percentage of time investigating the unfamiliar animal compared to the control (*Tsc1^flox/flox^*/Cre-negative) cohort. In this analysis, animals with targeted deletion of one *Tsc1* allele (*Tsc1^mut/wt^*) were not significantly different from *Tsc1*^mut/mut^ or control animal cohorts. Similar to the results from the elevated balance beam assay, within each tested genotype (*Tsc1*^mut/mut^, *Tsc1*^mut/wt^, and control), we found no significant difference in these values based on sex (P=0.2681, two-way ANOVA, Supplemental Table S1).

**Figure 4:**
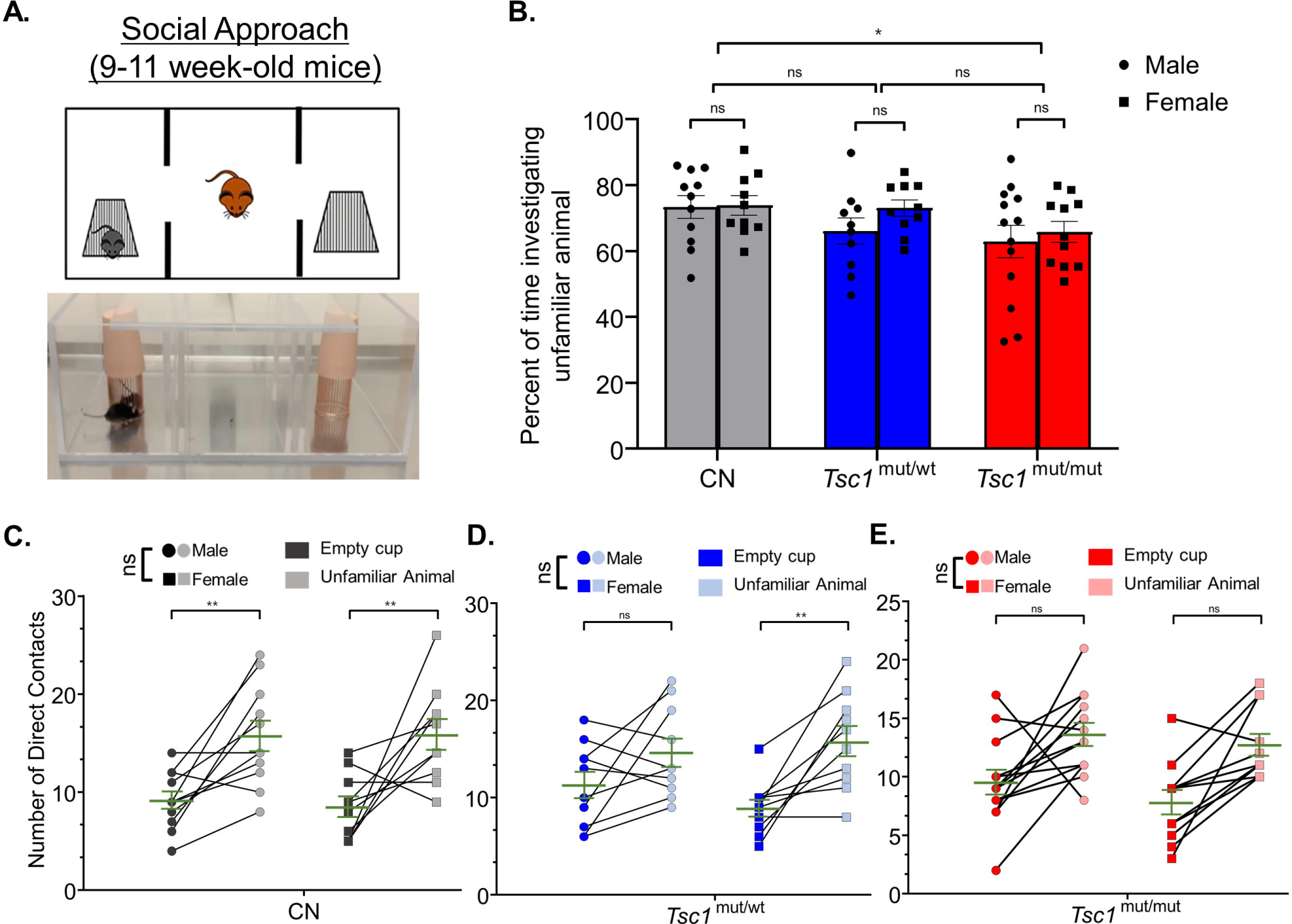
Social approach deficits are similar across males and females in control and *Tsc1* mutant animals. 9-11-week-old male and female *Tsc1*^mut/mut^ (N=13 males and N=11 females), *Tsc1*^mut/wt^ (N=10 males and N=10 females), and control (CN, N=11 males and N=10 females) mice were tested for social approach preference using the three-chamber apparatus depicted in in panel **A.** Photograph and cartoon depictions show the test animal, unfamiliar stranger mouse (in the left chamber), and empty wire cup (in the right chamber). **B.** *Tsc1*^mut/mut^ mice spent a significantly (p<0.05, two-way ANOVA) smaller proportion of time investigating the unfamiliar animal compared to *Tsc1*^mut/wt^ and control mice. No differences were recorded between sexes. In panels **C**, **D**, and **E**, the number of contacts with the empty cup and the unfamiliar animal is plotted for each test mouse for males and females of each genotype. Within each genotype, no significant difference was measured across sex for preference for contact with the unfamiliar animal. Individual values are plotted with the mean ± SEM shown in green (ns, not significant, * P< 0.05, ** P<0.01, Šídák’s corrected multiple comparisons tests used in two-way ANOVAs).

As another method to quantify the performance on the social approach behavioral test, the number of the test mouse’s direct contacts with the cage holding the unfamiliar animal and the number of direct contacts with the empty cage are plotted in Figure 4C, D, E, across sexes, for *Tsc1^mut/mut^*, *Tsc1*^mut/wt^, and control genotypes, respectively. These data reveal control mice of both sexes exhibit social approach preference (P<0.01, two-way ANOVA, Figure 4C), having a higher number of direct contacts with the unfamiliar animal compared to the empty cage. Interestingly, while male *Tsc1*^mut/mut^ and *Tsc1*^mut/wt^ mice did not exhibit significant preference for the unfamiliar animal over the empty cage (*Tsc1*^mut/mut^: P=0.1627, *Tsc1*^mut/wt^: P=0.1065, two-way ANOVA, Figure 4D, E), indicating impaired social approach behavior in these genotypes, female *Tsc1*^mut/wt^ mice did exhibit significant preference for the unfamiliar animal (P<0.01, two-way ANOVA, Figure 4D) similar to control animals; a phenotype not shared by *Tsc1*^mut/wt^ males.

In older (16-24 week-old) animals, however, the social approach testing paradigm revealed a clear difference between sexes in *Tsc1*^mut/mut^ mice (Fig. 5A, B). Male *Tsc1*^mut/mut^ mice spent a significantly smaller proportion of time than female test animals investigating the unfamiliar mouse (mean difference: −18.43±5.1, P<0.01, N=8 males and N=9 females, two-way ANOVA, Fig. 5B). No difference between sexes was measured in control mice (P=0.99, N=8 males and N=6 females, two-way ANOVA, Fig. 5B). In figure 5C, D, the number of the test mouse’s direct contacts with the cage holding the unfamiliar animal versus the empty cage are plotted between sexes for control and *Tsc1^mut/mut^* animals, respectively. Consistent with the results presented in Fig. 5B, control test mice from both sexes exhibited a significant preference for contacts with the unfamiliar animal compared to the empty cage (P=<0.0001, two-way ANOVA, Fig. 5C). In control mice, we measured no significant difference in preference between males and females (P=0.1123, two-way ANOVA, Fig. 5C). Conversely, male *Tsc1*^mut/mut^ mice exhibited a significantly (P=0.0117, two-way ANOVA, Fig. 5D) lesser preference for contacting the unfamiliar animal compared to female *Tsc1*^mut/mut^ mice.

**Figure 5:**
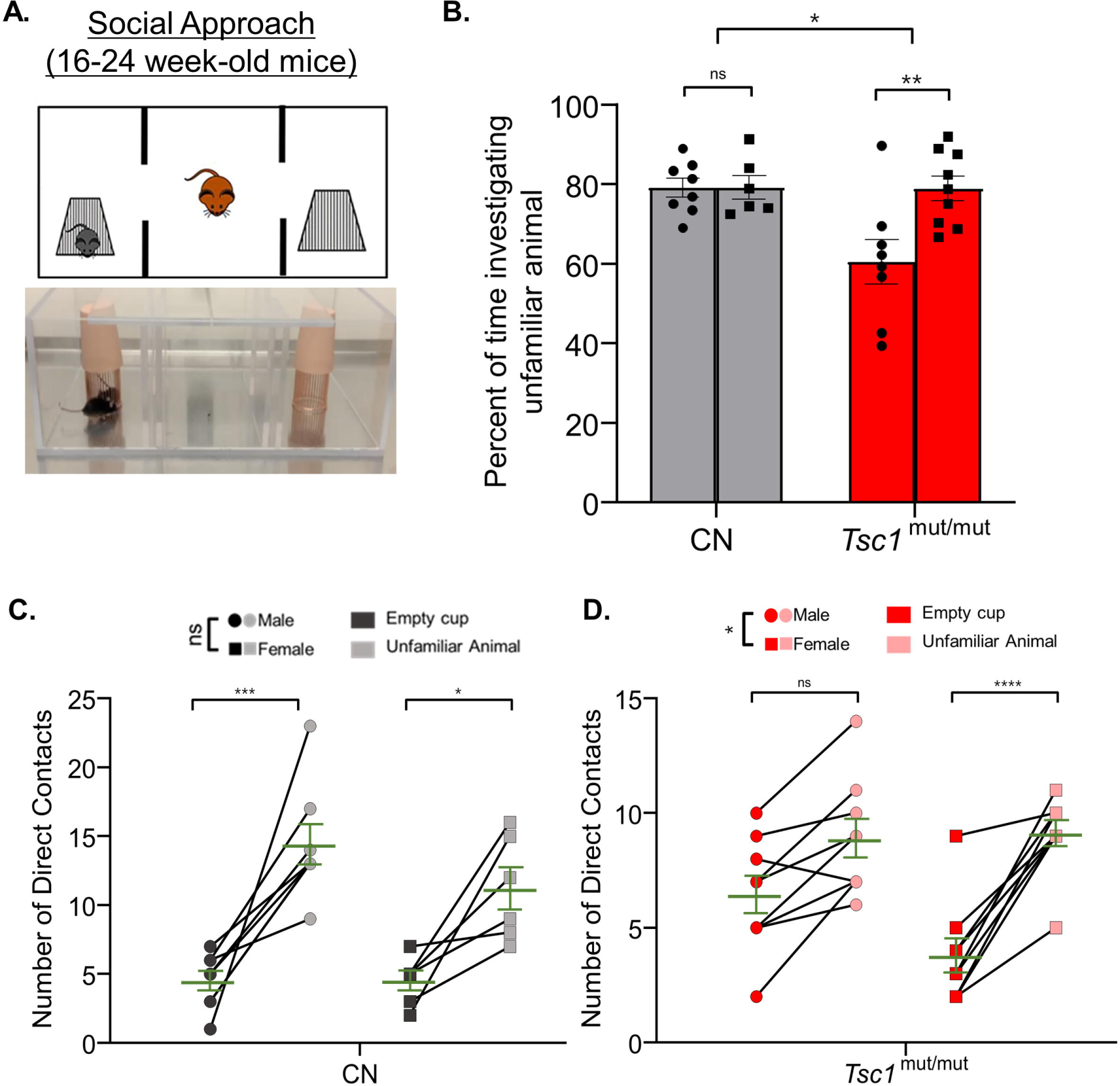
In 16-24 week-old *Tsc1^mut/mut^*mice, social approach deficits are more severe in males compared to females. 16-24 week-old male and female *Tsc1*^mut/mut^ (N=8 males and N=9 females) and control (CN, N=8 males and N=6 females) mice were tested using the three-chamber apparatus for social approach preference depicted in panel **A. B.** *Tsc1*^mut/mut^ mice spent a significantly (p<0.05, two-way ANOVA) lesser percentage of time investigating the unfamiliar animal compared to controls. Male *Tsc1*^mut/mut^ mice spent a significantly (p<0.05, two-way ANOVA) lesser percentage of time investigating the unfamiliar animal compared to *Tsc1^mut/mut^* females. In panels **C**, and **D**, the number of contacts with the empty cup and the unfamiliar animal is plotted for male and female test mice of control and *Tsc1^mut/mut^* genotypes. Male *Tsc1*^mut/mut^ mice showed a significantly (p<0.05, two-way ANOVA) lesser preference for contact with the unfamiliar animal than *Tsc1^mut/mut^* females. No difference between sexes was recorded in the control group. Individual values are plotted with the mean ± SEM shown in green (ns, not significant, * P< .05, ** P<0.01, *** P<0.001; **** P<0.0001, Šídák’s corrected multiple comparison tests used in two-way ANOVAs).

### 3.4 Sex specific differences in social approach behaviors were identified in both control and *Tsc1^mut/mut^* mice

To test for a preference for social novelty across sexes, the three-chamber apparatus was used to measure the preference of test animals for investigating an unfamiliar animal (in one chamber) compared to a familiar animal (in the alternative chamber). An image and cartoon depiction of this testing paradigm is presented in Figure 6A. Social novelty preference is measured as the preference for investigating the unfamiliar animal compared to the familiar animal. These tests were performed directly after social approach testing (see Methods). Also similar to our previous behavioral tests, in 9-11 week-old animals, a cohort of *Tsc1*^mut/wt^ animals was also tested. Interestingly, in these social novelty preference tests, we measured no significant difference in preference for investigating the unfamiliar animal across (9-11 week-old) control, *Tsc1*^mut/wt^, and *Tsc1*^mut/mut^ genotypes (P=0.2972, two-way ANOVA, Fig. 6B). This is in contrast to previous findings in which *Tsc1* mutant mice were found to exhibit no preference for investigating the socially novel animal (Tsai et al., 2012). Interestingly, within each genotype, control and *Tsc1*^mut/wt^ genotypes were found to have significant differences in social novelty preference based on sex. For instance, in control mice, female test animals investigated the unfamiliar animal a significantly lower percentage of time compared to male test animals (mean difference: 18.9 ± 5.8, P=0.0055, two-way ANOVA, Fig. 6B). These differences in preference based on sex were not, however, measured in the *Tsc1*^mut/mut^ cohort (P=0.9986, two-way ANOVA, Fig. 6B).

**Figure 6:**
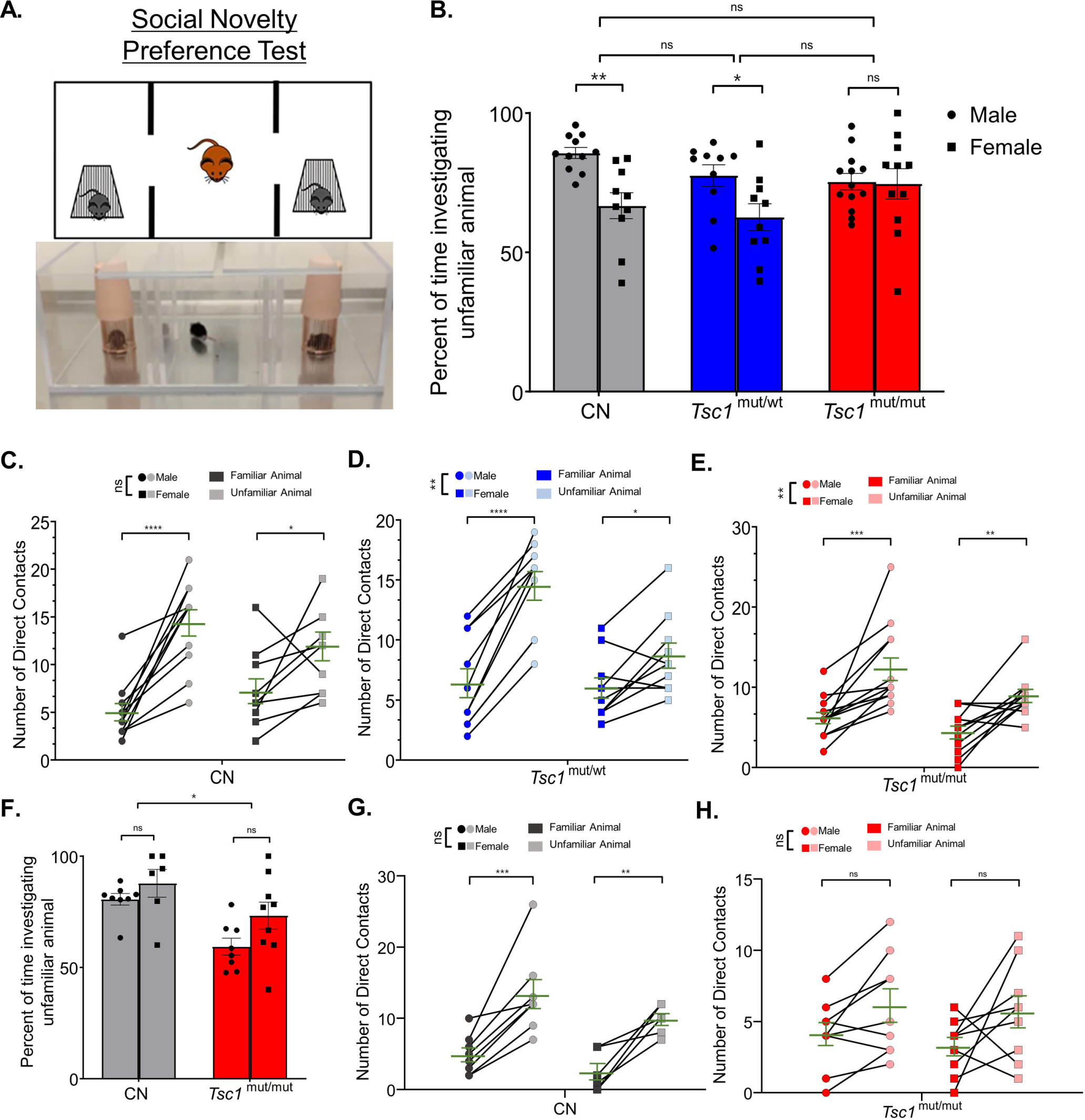
Sex-specific differences in social novelty preference occur in *Tsc1* mutant and control groups. 9-11-week-old male and female *Tsc1*^mut/mut^ (N=13 males and N=11 females), *Tsc1*^mut/wt^ (N=10 males and N=10 females), and control (CN, N=11 males and N=10 females) mice were tested for preference for social novelty using the three-chamber apparatus depicted in panel **A.** In the photograph and cartoon depiction, the test mouse in the center chamber is free to interact with an unfamiliar or familiar animal in the two side chambers. Panels **B-E.** show results for 9-11 week-old animal testing and panels **F-H.** show results for 16-24 week-old animal tests. **B.** Female control and *Tsc1*^mut/wt^ mice investigated the unfamiliar animal a significantly lesser percentage of time (CN: p<0.001, two-way ANOVA, *Tsc1*^mut/wt^: p<0.01, two-way ANOVA) compared to males. No significant differences were measured between genotypes. **C.** For the *Tsc1*^mut/mut^ and *Tsc1*^mut/wt^ groups, females had significantly (p<0.01, two-way ANOVA) lower preference for direct contact with the unfamiliar animal compared to males. Mean ± SEM values are plotted in **B.** In panel **C.**, individual values are plotted with the mean ± SEM shown in green (ns, not significant, * P<0.05, ** P<0.01, *** P<0.001; **** P<0.0001, Šídák’s corrected multiple comparison tests used in two-way ANOVAs). **F.** In 16-24 week-old *Tsc1*^mut/mut^ (N=8 males and N=9 females) and control (N=8 males and N=6 females) cohorts, *Tsc1*^mut/mut^ mice spent significantly (p<0.05, two-way ANOVA) less time investigating the unfamiliar mouse than controls. **G-H.** *Tsc1*^mut/mut^ mice had no preference for direct contact with the familiar mouse. No differences between sex were measured in control (**G.**) or *Tsc1^mut/mut^* (**H.**) animals. Mean ± SEM values are plotted in **F.** In panel **G-H.**, individual values are plotted with the mean ± SEM shown in green (ns, not significant, * P< 0.05, ** P<0.01, *** P<0.001, Šídák’s corrected multiple comparison tests used in two-way ANOVAs).

Interestingly, when number of cage contacts was used to assess social novelty preference in the 9-11 week-old animals, we measured no sex-specific differences in control animals (P=0.9222, two-way ANOVA, Fig. 6C), however, in *Tsc1^mut/wt^* (Fig. 6D) and *Tsc1^mut/mut^* (Fig. 6E) cohorts, there were significant differences based on sex, with females in both *Tsc1^mut/wt^* and *Tsc1^mut/mut^* cohorts contacting the unfamiliar animal cage less than males of the respective cohort (Fig. 6D, E). Together, these data suggest female test animals, from all genotypes tested, exhibit a lower preference for social novelty compared to male test animals; importantly, this sex-specific difference was measured in control animals, which makes assessing sex-specific effects of *Tsc1* deletion difficult. This difference between sexes was not found to be present in the *Tsc1*^mut/mut^ group. A previous study which compared social behavioral phenotypes of prairie voles to C57BL/6 mice also identified sex-specific differences in wild type C57BL/6 mice social novelty preference (Wu et al. 2022). Similar to the results presented here, Wu et al. (2022) determined male mice exhibit significantly greater social novelty preference (Fig. 6B) compared to female mice.

Social novelty performance was assessed again in 16-24 week old control and *Tsc1^mut/mut^*cohorts, with results presented in Fig. 6F, G and H. 16-24 week-old *Tsc1*^mut/mut^ mice spent a significantly smaller percentage of time investigating the unfamiliar animal than control mice (P=0.0438, two-way ANOVA, Fig. 6F), revealing a deficit in typical (control) social novelty preference. No significant differences between male and female test animals of either genotype were measured (control-P=0.5644, *Tsc1*^mut/mut^-P=0.0917, two-way ANOVA, Fig. 6G, H). Male and female control mice exhibited a significant preference for contacting the socially novel animal (P<0.0001, two-way ANOVA, Figure 6G), and, in contrast to the results from the 9-11 week-old cohorts, *Tsc1*^mut/mut^ mice did not exhibit preference for the unfamiliar mouse based on contact number (males: P=0.1889, females: 0.0942, two-way ANOVA, Fig. 6H). These data suggest older (16-24 week old) *Tsc1*^mut/mut^ develop an impairment in social novelty preference that is not significant at younger (9-11 week-old) ages.

## 4. DISCUSSION

Studies investigating sexual dimorphism in ASD symptoms in humans often result in inconclusive or contradictory findings. For example, there are reported sex differences in repetitive behaviors (Tillmann et al., 2018) and in fine motor control and social communication abilities (Craig et al., 2020), however, other investigations found no significant differences in these symptom domains across sexes (Andersson et al., 2013, Reinhardt et al., 2015). This emphasizes the need for a more targeted study of ASD symptomatology across sexes in animal models in which the underlying cell- and circuit-level drivers of behavioral phenotypes are well-established. This use of animal models to investigate ASD sex differences has only recently garnered increased interest; previous preclinical animal studies often only used males to avoid physiological variability linked to the estrous cycle of female rodents (Jeon et al. 2018). Interestingly, however, a study on of the effect of the female estrous cycle on behavioral testing, including social interaction assays, found very few parameters with significant differences between estrus and diestrus female mice (Zeng et al. 2023).

Here, we tested if the selective deletion of *Tsc1* in cerebellar Purkinje neurons differentially affects Purkinje neuron intrinsic excitability, as well as ASD-related behaviors in male and female mice. We tested animals in two (9-11 week-old and 16-24 week-old) age groups for social interaction and balance and motor coordination behaviors. Results from these studies revealed more severe deficits in older (16-24 week-old) male, compared to female, *Tsc1^mut/mut^* mice. This is similar to the more severe social interaction deficits identified in males (compared to females) of a *Shank3* mutant mouse model of ASD (Matas et al. 2021), although this mutant model relied on a global mutation, making it difficult to confidently attribute the sexual dimorphism to an effect on Purkinje neurons. Interestingly, we measured no significant differences between *Tsc1^mut/mut^* males and females on the elevated balance beam assay in both age groups, suggesting Purkinje neuron-specific *Tsc1* deletion affects cerebellar circuits that contribute to somatosensory function similarly across males and females; and conversely, in cerebellar circuits that contribute to social interaction behaviors, the selective deletion of *Tsc1* has a more severe effect in males compared to females.

### 4.1 Cell and circuit etiology of*Tsc1^mut/mut^* behavioral impairments

Numerous ASD-linked behavioral deficits were originally identified in (male) *Tsc1^mut/mut^*mice and these phenotypes were linked to two types of cerebellar impairment. In older (>10 week-old) *Tsc1^mut/mut^* animals, Tsai et al. (2012) measured reduced numbers of Purkinje neurons in the cerebellar vermis, as well as a significant increase in the proportion of Purkinje neurons expressing apoptotic factors TUNEL and Caspase 9. No changes in Purkinje neuron number were measured in younger (6 week-old) animals, however, there were marked deficits in Purkinje neuron repetitive firing properties (Tsai et al. 2012), similar to the results reported here (Figure 2). Because impaired behaviors were also recorded in younger (∼6 week-old animals) it’s likely that *Tsc1* deletion causes cerebellar circuit deficits through both impaired Purkinje neuron excitability, and, as the animals become older, by way of Purkinje neuron apoptosis. The stand-out result from the current investigation is that in older *Tsc1^mut/mut^*animals, impaired social interaction behavior (social approach preference) is more severe in *Tsc1^mut/mut^* males compared to females, and importantly, this sex-specific difference is not present in behaviors related to balance and motor coordination. This result indicates the sex-specific impairments vary across distinct cerebellar circuits. Previous work has also suggested the selective deletion of *Tsc1* from Purkinje neurons differentially affects distinct cerebellar circuits. Rapamycin treatment, which has been shown to rescue deficits associated with *Tsc1* deletion (Meikle et al. 2008; Zhou et al. 2009), was administered at various ages of *Tsc1^mut/mut^*mice to determine at which ages the treatment was effective in rescuing behavioral impairments. Interestingly, the ages at which rapamycin was effective at alleviating behavioral impairments were dependent on the type of behavior being rescued. For instance, rapamycin treatment in 10 week-old animals rescued deficits in motor function (measured via the rotarod test), but deficits in social interaction behaviors were not improved (Tsai et al. 2018). A clear hypothesis for this difference in rescue is that rapamycin-mediated rescue is no longer possible if Purkinje neurons in the relevant cerebellar circuit have undergone apoptosis. Extended to the work presented here, it may be that in male *Tsc1^mut/mut^* animals, there is greater apoptosis and Purkinje neuron loss in circuits contributing to social interaction behaviors compared to *Tsc1^mut/mut^* females, and that this sex-specific difference does not occur in cerebellar circuits contributing to balance and motor coordination. It will be interesting to test these ideas directly. Purkinje neurons functional in circuits dedicated to balance and motor coordination have been shown to localize to spinocerebellar (vermis and paravermis) regions (Unverdi and Alsayouri 2024); while Purkinje neurons functional in higher order behaviors, such as social interactions have been associated with Purkinje neurons localized in hemispheric Crus I and Crus II cerebellar subregions (Matas et al. 2021; Stoodley et al. 2017; Van Overwalle et al. 2020). Of course, in addition, or perhaps in place of sex-specific differences in Purkinje neuron death, there may be sex-specific and circuit specific differences in the deficits in Purkinje neuron membrane excitability of *Tsc1^mut/mut^* animals, which should also be tested. Finally, it’s of note that the sex-specific difference in social approach behaviors of *Tsc1^mut/mut^* animals was not measured in younger (9-11 week-old) animals. This result is in line with the progressive nature of behavioral deficits in *Tsc1^mut/mut^* animals (Tsai et al. 2018; 2012), which also may result from progressive Purkinje neuron apoptosis that occurs differentially across distinct cerebellar circuits. Beyond identifying the cellular and circuit-level substrates responsible for the sexual dimorphism in this ASD model, it will be interesting to also investigate the physiological drivers of such differences. For instance, there is some evidence that female sex hormones may provide a neuroprotective function in neurodegenerative disorders (Sumien et al. 2021; Túnez et al. 2006; Henderson et al. 1994; Garcia-Segura, Azcoitia, and DonCarlos 2001), resulting in sex-specific outcomes, although differences in the expression properties of sex chromosomes and/or sex hormone receptor expression (Ng and Hazrati 2022) may also contribute to these differences. In future work aimed at effectively linking *Tsc1^mut/mut^*behavioral impairments with impairments in cerebellar functioning, it will be necessary to categorize experiments based on animal sex and age, and in cellular-based experiments, careful localization and targeting of Purkinje neurons should be used to link the functioning of discrete cerebellar circuits with the relevant behavioral phenotype.

## 6. AUTHOR CONTRIBUTIONS

RJL and JLR planned experiments; RJL and NJL performed and analyzed the results of behavioral experiments; SPB performed and analyzed the results of electrophysiology experiments; AKJ and JJO performed microscopy experiments; RJL and JLR wrote the manuscript. All authors reviewed and approved the manuscript prior to submission.

## 8. ACKNOWLEDGMENTS

The authors gratefully acknowledge and thank personnel of the Miami University Laboratory Animal Resources (LAR) and Center for Advanced Microscopy and Imaging (CAMI).

