## Supplemental Tables and Figures for "Selective deletion of *Tsc1* from mouse cerebellar Purkinje neurons drives sex-specific behavioral impairments linked to autism"

**Supplemental Table 1: Results of statistical analyses for 9-11 week-old mice.**
Main effects of two-way ANOVA tests across sexes for each behavioral assay are reported as P-values. P<0.05 was considered to be statistically significant; significant effects are bolded. Dash indicates that a pairwise comparison was unable to be completed; RM, repeated measures.

| N | Test | Test Details | *Post hoc* Multiple Comparisons |
|---|---|---|---|
| **Elevated Balance Beam** | | | |
| N=21<br>Males=11<br>Females = 10 | CN Males & Females<br>Time to Cross (10mm):<br>RM 2-way ANOVA | day x sex: F(3,57)=0.8883, P=0.4528<br>sex: F(1,19)=0.3849, P=0.5424<br>day: F(2.592,49.24)=0.8088, P=0.4794 | Šídák's, Male vs. Female<br>Day 1: P=0.5957<br>Day 2: P=0.9999<br>Day 3: P=0.9954<br>Day 4: P=0.8159 |
| N=20<br>Males=10<br>Females = 10 | *Tsc1*$^{mut/wt}$ Males & Females<br>Time to Cross (10mm):<br>RM 2-way ANOVA | sex x day: F(3,54)=0.1186, P=0.9488<br>sex: F(1,18)=1.784, P=0.1983<br>day: F(2.078,37.41)=1.630, P=0.2089 | Šídák's, Male vs. Female<br>Day 1: P=0.8600<br>Day 2: P=0.9072<br>Day 3: P=0.5291<br>Day 4: P=0.7677 |
| N=24<br>Males=13<br>Females = 11 | *Tsc1*$^{mut/mut}$ Males & Females<br>Time to Cross (10mm):<br>RM 2-way ANOVA | sex x day: F(3,66)=1.882, P=0.1412<br>sex: F(1,22)=2.828, P=0.1068<br>day: F(2.175,47.86)=1.630, P=0.2089 | Šídák's, Male vs. Female<br>Day 1: P=0.1941<br>Day 2: P=0.1873<br>Day 3: P=0.8345<br>Day 4: P=0.9753 |
| N=21<br>Males=11<br>Females = 10 | CN Males & Females<br>Time to Cross (8mm):<br>RM 2-way ANOVA | sex x day: F(3,57)=0.08166, P=0.9697<br>sex: F(1,19)=4.365, P=0.0504<br>day: F(2.530,48.07)=0.1666, P=0.8912 | Šídák's, Male vs. Female<br>Day 1: P=0.6696<br>Day 2: P=0.1831<br>Day 3: P=0.7248<br>Day 4: P=0.6880 |
| N=20<br>Males=10<br>Females = 10 | *Tsc1*$^{mut/wt}$ Males & Females<br>Time to Cross (8mm):<br>RM 2-way ANOVA | sex x day: F(3,54)=0.028667, P=0.9940<br>sex: F(1,18)=1.016, P=0.3269<br>day: F(2.805,50.49)=1.372, P=0.2627 | Šídák's, Male vs. Female<br>Day 1: P=0.4348<br>Day 2: P=0.9786<br>Day 3: P=0.7831<br>Day 4: P=0.9490 |
| N=24<br>Males=13<br>Females = 11 | *Tsc1*$^{mut/mut}$ Males & Females<br>Time to Cross (8mm):<br>RM 2-way ANOVA | sex x day: F(3,66)=1.073, P=0.3667<br>sex: F(1,22)=0.04588, P=0.8324<br>day: F(1.938,42.63)=1.330, P=0.2747 | Šídák's, Male vs. Female<br>Day 1: P=0.9623<br>Day 2: P=0.9119<br>Day 3: P=0.9853<br>Day 4: P=0.9845 |
| N=21<br>Males=11<br>Females = 10 | CN Males & Females<br>Footslips (10mm):<br>RM 2-way ANOVA | sex x day: F(3,57)=2.184, P=0.0986<br>sex: F(1,19)=0.3501, P=0.5610<br>day: F(2.029,38.55)=0.9767, P=0.3867 | Šídák's, Male vs. Female<br>Day 1: P=0.8113<br>Day 2: P>0.9999<br>Day 3: P=0.5205<br>Day 4: -- |

| | | | |
|---|---|---|---|
| N=20<br>Males=10<br>Females = 10 | *Tsc1*$^{mut/wt}$ Males & Females Footslips (10mm):<br>RM 2-way ANOVA | sex x day: F(3,54)=0.8319, P=0.4822<br>sex: F(1,18)=1.528, P=0.2323<br>day: F(2.201,39.62)=0.8319, P=0.4527 | Šídák's, Male vs. Female<br>Day 1: P>0.9999<br>Day 2: P=0.5205<br>Day 3: --<br>Day 4: P=0.8142 |
| N=24<br>Males=13<br>Females = 11 | *Tsc1*$^{mut/mut}$ Males & Females Footslips (10mm):<br>RM 2-way ANOVA | sex x day: F(3,66)=0.3565, P=0.7846<br>sex: F(1,22)=0.1423, P=0.7056<br>day: F(2.591,57)=2.834, P=0.0536 | Šídák's, Male vs. Female<br>Day 1: P>0.9999<br>Day 2: P=0.9607<br>Day 3: P=0.9998<br>Day 4: P=0.9061 |
| N=21<br>Males=11<br>Females = 10 | CN Males & Females Footslips (8mm):<br>RM 2-way ANOVA | sex x day: F(3,57)=1.182, P=0.3248<br>sex: F(1,19)=0.3073, P=0.5858<br>day: F(2.557,48.59)=0.9318, P=0.4204 | Šídák's, Male vs. Female<br>Day 1: P=0.5953<br>Day 2: P=0.7766<br>Day 3: P=0.9931<br>Day 4: P=0.6969 |
| N=20<br>Males=10<br>Females = 10 | *Tsc1*$^{mut/wt}$ Males & Females Footslips (8mm):<br>RM 2-way ANOVA | sex x day: F(3,54)=0.9649, P=0.4160<br>sex: F(1,18)=1.477, P=0.2400<br>day: F(2.6,46.8)=1.076, P=0.3621 | Šídák's, Male vs. Female<br>Day 1: P=0.9981<br>Day 2: P=0.9960<br>Day 3: P=0.1967<br>Day 4: P=0.9090 |
| N=24<br>Males=13<br>Females = 11 | *Tsc1*$^{mut/mut}$ Males & Females Footslips (8mm):<br>RM 2-way ANOVA | sex x day: F(3,66)=1.160, P=0.3318<br>sex: F(1,22)=0.02369, P=0.8791<br>day: F(2.539,55.86)=5.157, **P=0.0051** | Šídák's, Male vs. Female<br>Day 1: P=0.9985<br>Day 2: P=0.8292<br>Day 3: P=0.9969<br>Day 4: P=0.8586 |
| | | **Social Interaction** | |
| N=21<br>Males=11<br>Females = 10 | CN Males & Females Direct Contacts (Approach):<br>RM 2-way ANOVA | sex x preference: F(1,19)=0.08960, P=0.7679<br>sex: F(1,19)=0.04783, P=0.8292<br>preference: F(1,19)=23.86, **P=0.0001** | Šídák's, Empty cup vs. unfamiliar animal<br>Male: **P=0.0071**<br>Female: **P=0.0040** |
| N=20<br>Males=10<br>Females = 10 | *Tsc1*$^{mut/wt}$ Males & Females Direct Contacts (Approach):<br>RM 2-way ANOVA | sex x preference: F(1,18)=1.994, P=0.1750<br>sex: F(1,18)=0.1828, P=0.6741<br>preference: F(1,18)=16.01, **P=0.0008** | Šídák's, Empty cup vs. unfamiliar animal<br>Male: P=0.1605<br>Female: **P=0.0025** |
| N=24<br>Males=13<br>Females = 11 | *Tsc1*$^{mut/mut}$ Males & Females Direct Contacts (Approach):<br>RM 2-way ANOVA | sex x preference: F(1,22)=1.639, P=0.6895<br>sex: F(1,22)=1.694, P=0.2065<br>preference: F(1,22)=19.11, **P=0.002** | Šídák's, Empty cup vs. unfamiliar animal<br>Male: P=0.1627<br>Female: P=0.0845 |

| | | | |
|---|---|---|---|
| N=21<br>Males=11<br>Females = 10 | CN Males & Females Direct Contacts (Novelty):<br>RM 2-way ANOVA | sex x preference: $F(1,19)=3.604$, $P=0.0729$<br>sex: $F(1,19)=0.009826$, $P=0.9222$<br>preference: $F(1,19)=32.77$, **$P<0.0001$** | Šídák's, Familiar animal vs. unfamiliar animal<br>Male: **$P<0.0001$**<br>Female: **$P=0.0318$** |
| N=20<br>Males=10<br>Females = 10 | *Tsc1*$^{mut/wt}$ Males & Females Direct Contacts (Novelty):<br>RM 2-way ANOVA | sex x preference: $F(1,18)=17.61$, **$P=0.0005$**<br>sex: $F(1,18)=4.992$, **$P=0.0384$**<br>preference: $F(1,18)=70.45$, **$P<0.0001$** | Šídák's, Familiar animal vs. unfamiliar animal<br>Male: **$P<0.0001$**<br>Female: **$P=0.0164$** |
| N=24<br>Males=13<br>Females = 11 | *Tsc1*$^{mut/mut}$ Males & Females Direct Contacts (Novelty):<br>RM 2-way ANOVA | sex x preference: $F(1,22)=0.6437$, $P=0.4310$<br>sex: $F(1,22)=5.774$, **$P=0.0251$**<br>preference: $F(1,22)=30.97$, **$P<0.0001$** | Šídák's, Familiar animal vs. unfamiliar animal<br>Males: **$P=0.0002$**<br>Females: **$P=0.0076$** |
| N=65<br>CN=11 Males, 10 Females<br>*Tsc1*$^{mut/wt}$ = 10 Males, 10 Females<br>*Tsc1*$^{mut/muut}$ = 13 Males, 11 Females | CN, *Tsc1*$^{mut/wt}$, & *Tsc1*$^{mut/mut}$, Males & Females<br>Percentage of Investigation (Approach): 2-way ANOVA | sex x genotype: $F(2,59)=0.3772$, $P=0.6874$<br>genotype: $F(2,59)=3.220$, **$P=0.0471$**<br>sex: $F(1,59)=1.250$, $P=0.2681$ | Šídák's, Genotype<br>CN vs. *Tsc1*$^{mut/wt}$: $P=0.6523$<br>CN vs. *Tsc1*$^{mut/mut}$: **$P=0.0424$**<br>*Tsc1*$^{mut/wt}$ vs. *Tsc1*$^{mut/mut}$: $P=0.4173$ |
| N=65<br>CN=11 Males, 10 Females<br>*Tsc1*$^{mut/wt}$ = 10 Males, 10 Females<br>*Tsc1*$^{mut/muut}$ = 13 Males, 11 Females | CN, *Tsc1*$^{mut/wt}$, & *Tsc1*$^{mut/mut}$, Males & Females<br>Percentage of Investigation (Novelty): 2-way ANOVA | sex x genotype: $F(2,59)=2.922$, $P=0.0617$<br>genotype: $F(2,59)=1.239$, $P=0.2972$<br>sex: $F(1,59)=12.18$, **$P=0.0009$** | Šídák's, Male vs. Female<br>CN: **$P=0.0055$**<br>*Tsc1*$^{mut/wt}$: **$P=0.0434$**<br>*Tsc1*$^{mut/mut}$: $P=0.9986$ |
| **Repetitive Behavior** | | | |
| N=43<br>CN=6 Males, 8 Females<br>*Tsc1*$^{mut/wt}$ = 6 Males, 5 Females<br>*Tsc1*$^{mut/muut}$ = 9 Males, 9 Females | CN, *Tsc1*$^{mut/wt}$, & *Tsc1*$^{mut/mut}$, Males & Females<br>Total Time Grooming:<br>2-way ANOVA | sex x genotype: $F(2,37)=2.414$, $P=0.1034$<br>genotype: $F(2,37)=0.2444$, $P=0.7844$<br>sex: $F(1,37)=4.682$, **$P=0.0370$** | Šídák's, Male vs. Female<br>CN: $P=0.0500$<br>*Tsc1*$^{mut/wt}$: $P=0.4180$<br>*Tsc1*$^{mut/mut}$: $P=0.9752$ |

**Supplemental Table 2: Results of statistical analyses for 16-24 week-old mice.**
Main effects of two-way ANOVA tests across sexes for each behavioral assay are reported as P-values. P <0.05 was considered to be statistically significant; significant effects are bolded. Dash indicates that a pairwise comparison was unable to be completed; RM, repeated measures.

| N | Test | Test Details | *Post hoc* Multiple Comparisons |
|---|---|---|---|
| | | Elevated Balance Beam | |
| N=18<br>Males=10<br>Females=8 | CN Males & Females Time to Cross (10mm):<br>RM 2-way ANOVA | sex x day: $F(3,48)=0.6286$, $P=0.6001$<br>sex: $F(1,16)=1.279$, $P=0.2930$<br>day: $F(2.703,43.25)=0.2806$, $P=0.6036$ | Šídák's, Male vs. Female<br>Day 1: P=0.7256<br>Day 2: P= 0.9976<br>Day 3: P=0.9871<br>Day 4: P=0.9992 |
| N=19<br>Males=9<br>Males=10 | *Tsc1*$^{mut/mut}$ Males & Females Time to Cross (10mm):<br>RM 2-way ANOVA | sex x day: $F(3,51)=0.7325$, $P=0.5374$<br>sex: $F(1,17)=2.595$, $P=0.12556$<br>day: $F(2.896,49.24)=1.746$, $P=0.1714$ | Šídák's, Male vs. Female<br>Day 1: P=0.3835<br>Day 2: P=0.9159<br>Day 3: P=0.4280<br>Day 4: P=0.6606 |
| N=18<br>Males=10<br>Females=8 | CN Males & Females Time to Cross (8mm):<br>RM 2-way ANOVA | sex x day: $F(3,48)=0.77699$, $P=0.5126$<br>sex: $F(1,16)=0.2006$, $P=0.6602$<br>day: $F(2.719,43.5)=0.9122$, $P=0.4350$ | Šídák's, Male vs. Female<br>Day 1: P=0.9998<br>Day 2: P>0.9999<br>Day 3: P=0.6919<br>Day 4: P=0.9515 |
| N=19<br>Males=9<br>Males=10 | *Tsc1*$^{mut/mut}$ Males & Females Time to Cross (8mm):<br>RM 2-way ANOVA | sex x day: $F(3,51)=0.1847$, $P=0.9063$<br>sex: $F(1,17)=1.010$, $P=0.3290$<br>day: $F(2.012,34.21)=2.271$, $P=0.1183$ | Šídák's, Male vs. Female<br>Day 1: P=0.9525<br>Day 2: P=0.8574<br>Day 3: P=0.6724<br>Day 4: P=0.8931 |
| N=18<br>Males=10<br>Females=8 | CN Males & Females Footslips (10mm):<br>RM 2-way ANOVA | sex x day: $F(3,48)=1.183$, $P=0.3260$<br>sex: $F(1,16)=0.1616$, $P=0.6930$<br>day: $F(2.395,38.31)=0.3188$, $P=0.7668$ | Šídák's, Male vs. Female<br>Day 1: P=0.8222<br>Day 2: P=0.8142<br>Day 3: P=0.8142<br>Day 4: -- |
| N=19<br>Males=9<br>Males=10 | *Tsc1*$^{mut/mut}$ Males & Females Footslips (10mm):<br>RM 2-way ANOVA | sex x day: $F(3,51)=1.904$, $P=0.1406$<br>sex: $F(1,17)=0.04234$, $P=0.8394$<br>day: $F(2.526,42.94)=1.1055$, $P=0.3510$ | Šídák's, Male vs. Female<br>Day 1: P=0.9978<br>Day 2: P=0.9772<br>Day 3: P=0.8486<br>Day 4: P=0.8643 |
| N=18<br>Males=10<br>Females=8 | CN Males & Females Footslips (8mm):<br>RM 2-way ANOVA | sex x day: $F(3,48)=0.8063$, $P=0.4966$<br>sex: $F(1,16)=2.538$, $P=0.1307$<br>day: $F(2.618,41.89)=1.936$, $P=0.1454$ | Šídák's, Male vs. Female<br>Day 1: P=0.5121<br>Day 2: P=0.9507<br>Day 3: P=0.9081<br>Day 4: P=0.9998 |

| | | | |
|---|---|---|---|
| N=19<br>Males=9<br>Males=10 | *Tsc1*$^{mut/mut}$ Males & Females Footslips (8mm):<br>RM 2-way ANOVA | sex x day: F(3,51)=0.9218, P=0.4370<br>sex: F(1,17)=0.08584, P=0.7731<br>day: F(2.193,37.29)=1.280, P=0.2918 | Šídák's, Male vs. Female<br>Day 1: P=0.9249<br>Day 2: P=0.9642<br>Day 3: P=0.9911<br>Day 4: P=0.9995 |
| | | Social Interaction | |
| N=14<br>Males=8<br>Females=6 | CN Males & Females Direct Contacts (Approach):<br>RM 2-way ANOVA | sex x preference: F(1,12)=1.323, P=0.2725<br>sex: F(1,12)=2.937, P=0.1123<br>preference: F(1,12)=35.16, **P<0.0001** | Šídák's, Empty cup vs. unfamiliar animal<br>Male: **P=0.0003**<br>Female: **P=0.0163** |
| N=18<br>Males=9<br>Males=9 | *Tsc1*$^{mut/mut}$ Males & Females Direct Contacts (Approach):<br>RM 2-way ANOVA | sex x preference: F(1,16)=8.096, **P=0.0117**<br>sex: F(1,16)=1.729, P=0.2071<br>preference: F(1,16)=58.68, **P<0.0001** | Šídák's, Empty cup vs. unfamiliar animal<br>Male: P=0.5288<br>Female: **P=0.0025** |
| N=14<br>Males=8<br>Females=6 | CN Males & Females Direct Contacts (Novelty):<br>RM 2-way ANOVA | sex x preference: F(1,12)=0.2264, P=0.6427<br>sex: F(1,12)=3.252, P=0.0965<br>preference: F(1,12)=41.70, **P<0.0001** | Šídák's, Empty cup vs. unfamiliar animal<br>Male: **P=0.0004**<br>Female: **P=0.0038** |
| N=18<br>Males=9<br>Males=10 | *Tsc1*$^{mut/mut}$ Males & Females Direct Contacts (Novelty):<br>RM 2-way ANOVA | sex x preference: F(1,16)=8.096, **P=0.0117**<br>sex: F(1,16)=1.729, P=0.2071<br>preference: F(1,16)=7.556, **P=0.0143** | Šídák's, Empty cup vs. unfamiliar animal<br>Male: P=0.1889<br>Female: P=0.0942 |
| N=32<br>CN=8 Males, 6 Females<br>*Tsc1*$^{mut/mut}$=9 Males, 9 Females | CN & *Tsc1*$^{mut/mut}$ Males & Females Percentage of Investigation (Approach): 2-way ANOVA | sex x genotype: F(1,27)=5.713, **P=0.0241**<br>genotype: F(1,27)=6.065, **P=0.0205**<br>sex: F(1,27)=5.848, **P=0.0226** | Šídák's, Male vs. Female<br>CN: P=0.9998<br>*Tsc1*$^{mut/mut}$: **P=0.0026** |
| N=32<br>CN=8 Males, 6 Females<br>*Tsc1*$^{mut/mut}$=9 Males, 9 Females | CN & *Tsc1*$^{mut/mut}$ Males & Females Percentage of Investigation (Novelty): 2-way ANOVA | sex x genotype: F(1,27)=0.4497, P=0.5082<br>genotype: F(1,27)=12.92, **P=0.0013**<br>sex: F(1,27)=4.472, **P=0.0438** | Šídák's, Male vs. Female<br>CN: P=0.5644<br>*Tsc1*$^{mut/mut}$: P=0.0917 |

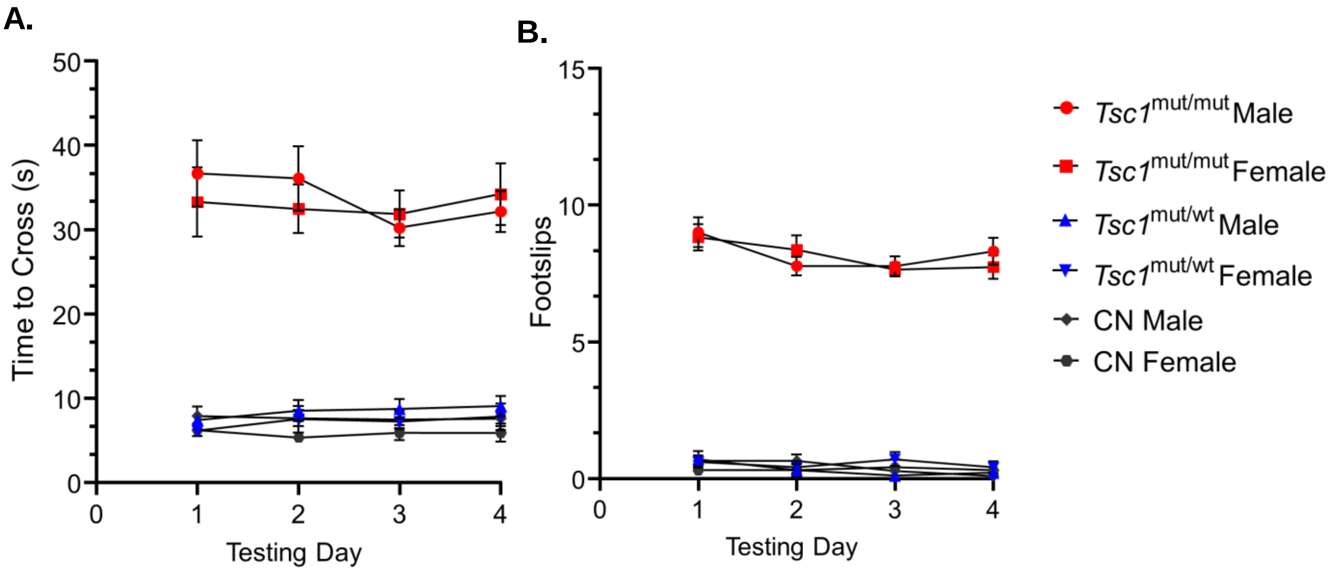


**Supplemental Figure 1**:

Motor coordination and balance testing of male and female 9–11 week-old *Tsc1*$^{mut/mut}$ (N=13 males and N=11 females), *Tsc1*$^{mut/wt}$ (N=10 males and N=10 females), and control (CN, N=11 males and N=10 females) mice were tested on an elevated balance beam 8mm in width. *Tsc1*$^{mut/mut}$ mice performed significantly ($p<0.0001$) worse than control animals on across all four testing days, but no difference in the time to cross (panel **A**) or number of hindlimb footslips (panel **B**) between sexes within each of the genotypes. Mean ± SEM values are plotted.

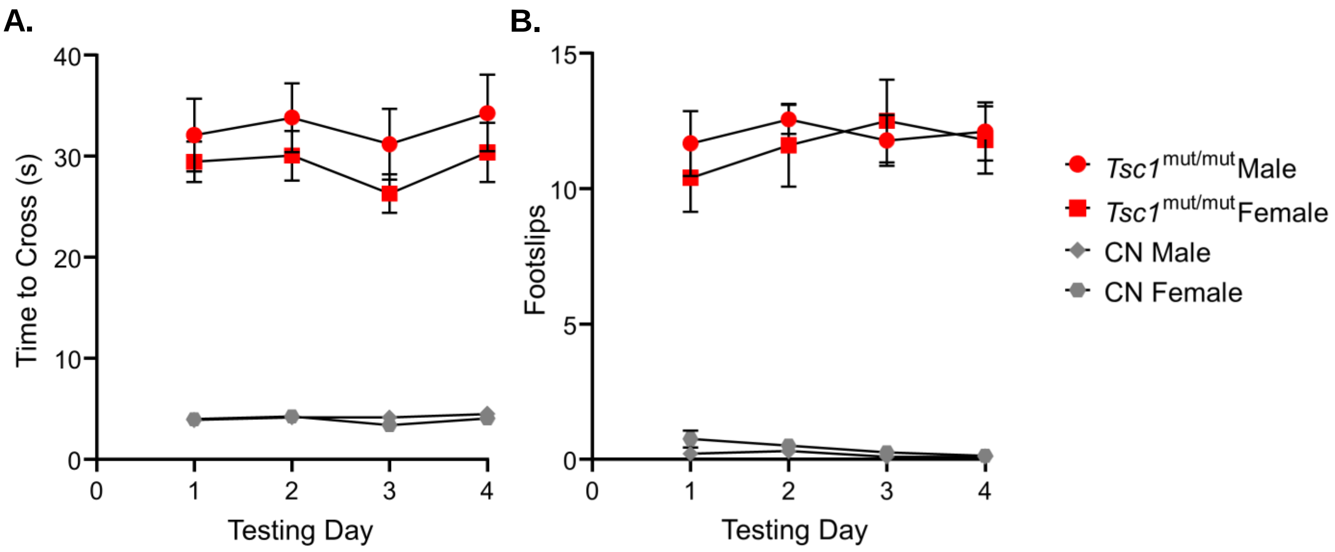


**Supplemental Figure 2.**
16-24 week-old *Tsc1*$^{mut/mut}$ (N=9 males and N=10 females) and control (N=10 males and N=8 females) mice of both sexes were tested on an elevated balance beam 8mm in width. *Tsc1*$^{mut/mut}$ mice performed significantly ($p<0.0001$) worse than control animals on across all four testing days, and no difference in the time to cross (panel **A**) or number of hindlimb footslips (panel **B**) between sexes was measured for either genotype. Mean ± SEM values are plotted.
